# Size-invariant exercise-induced aerobic scope across a broad body mass range in a giant isopod

**DOI:** 10.64898/2026.01.28.702439

**Authors:** Shogo Tanaka, Sayano Anzai, Shoma Izumi, Mitsuharu Yagi

**Affiliations:** Faculty of Fisheries, Nagasaki University, Nagasaki 852-8521, Japan

**Keywords:** aerobic scope, metabolic scaling, deep sea, scavenger, crustacean, Bathynomus

## Abstract

Aerobic scope is the aerobic metabolism left after maintenance costs are met, and it sets a simple limit on how much oxygen-dependent activity an animal can support. In many aquatic animals, this margin changes with body size because resting and active metabolic rates do not always scale in the same way. Large deep-sea scavengers are interesting in this respect because they live where food is sparse and unpredictable, but they must still move when carrion or food cues appear. We measured resting metabolic rate (RMR) and post-exercise peak oxygen consumption, here referred to as active metabolic rate (AMR), in the giant isopod *Bathynomus doederleini* across a body-mass range of 1.7‒48.4 g using intermittent-flow respirometry at 10 °C. RMR and AMR both increased with body mass and had similar scaling exponents. As a result, factorial aerobic scope (FAS = AMR/RMR) did not change with body mass (median FAS = 2.83). A residual analysis also showed that mass-corrected RMR was not significantly related to mass-corrected AMR. Thus, individuals with relatively high maintenance metabolism did not necessarily have higher post-exercise aerobic capacity. These results show that exercise-induced aerobic scope was maintained across ontogeny in this deep-sea scavenger. They also show that body size and individual metabolic phenotype provide different information: body size explained the shared increase in RMR and AMR, whereas residual variation did not show a strong link between the two traits.

## 1. Introduction

Aerobic scope represents the aerobic metabolic capacity available for biological functions beyond maintenance metabolism. It provides an important physiological link between whole-animal metabolism to ecological performance, indicating the capacity for activity, feeding, growth, reproduction, and recovery from disturbances (Fry, 1971; Claireaux and Lefrançois, 2007). In aquatic ectotherms, aerobic scope is typically estimated from the difference or ratio between resting or standard metabolic rate and the highest metabolic rate during activity or stimulation. Since body size strongly influences metabolic rate, understanding aerobic scope requires considering how resting and activity-related metabolic rates scale with body mass (Kraskura et al., 2023). Usually, both rates increase with body mass, but their relative scaling varies among species, ecological contexts, and physiological states (Brett, 1965; Glazier, 2005, 2009, 2010). Maximal metabolic rates are affected not only by maintenance costs but also by locomotor demands, oxygen transport, tissue metabolism, and ecological activity level (Jimenez et al., 2008; Jensen et al., 2013). Therefore, aerobic scope can increase, decrease, or stay constant during growth, depending on the scaling of resting versus activity-related metabolism. If activity-related metabolism increases more steeply, larger individuals may have a greater aerobic margin; if resting metabolism scales more steeply, they may have less capacity beyond maintenance requirements. Factorial aerobic scope is therefore useful for evaluating whether proportional aerobic capacity is maintained, reduced, or enhanced relative to absolute metabolic demand increases during growth (Killen et al., 2007; Zhang et al., 2014). Ontogenetic changes in aerobic scope have been studied most extensively in fishes. In some marine teleosts, resting and maximum metabolic rates scale differently during growth, producing mass-dependent changes in aerobic scope and suggesting that larger individuals may have “little left in the tank” (Killen et al., 2007). Other studies have shown that the scaling of metabolic rate can vary with lifestyle, temperature, and activity level, indicating that metabolic scaling is not governed by body size alone (Glazier, 2009, 2010; Zhang et al., 2014). These findings suggest that ontogenetic metabolic scaling should be interpreted in relation to life history and ecological context. Growth can alter locomotor demand, tissue composition, habitat use, and energy allocation, and these changes may affect the balance between maintenance metabolism and activity-related aerobic capacity (Post and Lee, 1996; Killen et al., 2007). This issue may be especially relevant to animals whose post-hatching development is not accompanied by major shifts in habitat or locomotor mode, because such species may maintain a relatively stable balance between maintenance demand and activity-related aerobic capacity across ontogeny.

Deep-sea scavengers provide a useful system for examining this issue. The deep sea is characterized by low temperature, darkness, high hydrostatic pressure, and limited or unpredictable food supply. In such environments, biological activity and resource availability can be spatially and temporally heterogeneous (Dayton and Hessler, 1972; Seibel and Drazen, 2007). Large organic falls and carrion can provide spatially localized and episodic resource pulses on the deep-sea floor (Smith and Baco, 2003). Benthic scavengers that depend on these resources may therefore experience extended periods of limited food availability interrupted by shorter periods of food detection, movement, feeding, digestion, and recovery. Under these conditions, a high resting metabolic rate may impose a continuous energetic cost, whereas high activity-related aerobic capacity may be beneficial mainly during episodic events. Seibel and Drazen (2007) emphasized that metabolic rates in marine animals are shaped not only by environmental constraints but also by ecological energy demand. The balance between resting and activity-related metabolism may therefore be particularly important in deep-sea scavengers, in which chronic food limitation coexists with the need for episodic activity.

The giant isopod *Bathynomus doederleini* is a benthic scavenger distributed in the northwestern Pacific (Lowry and Dempsey, 2006; Anzai et al., 2026). Recent ecological work has shown that its distribution and body-mass structure vary with depth, suggesting that body mass is closely related to habitat use in this species (Anzai et al., 2026). Metabolic responses to feeding and temperature have also been examined in *B. doederleini*, indicating that this species is a useful study system for investigating the physiology of deep-sea scavenging crustaceans (Tanaka et al., 2023). However, it remains unclear how resting and activity-related aerobic metabolism scale with body mass in *Bathynomus*, whether larger individuals maintain the same proportional aerobic capacity as smaller individuals, and whether the two metabolic traits covary among individuals after accounting for body mass. Residual-based analyses have been used to examine such body-mass-independent relationships in fishes, but comparable evidence remains scarce for large deep-sea crustaceans (Norin and Malte, 2011, 2012; Auer et al., 2017; Reemeyer and Rees, 2020).

Here, we examined the body mass scaling of resting metabolic rate (RMR) and post-exercise peak oxygen consumption, used here as active metabolic rate (AMR), in *B. doederleini* across a broad body-mass range. Specifically, we tested the scaling relationships of RMR and AMR, the size dependence of factorial aerobic scope (FAS = AMR/RMR), and the covariation between the two metabolic traits after accounting for body mass. By combining allometric and residual analyses, we evaluated whether proportional aerobic capacity is maintained across ontogeny and whether population-level metabolic scaling reflects individual-level covariation between resting and activity-related metabolism

## 2. Materials and methods

### 2.1 Animal collection and maintenance

Giant isopods *Bathynomus doederleini*, were collected between April 2024 and August 2025 at depths of 201‒764 m off the Goto Islands, Japan. Sampling was conducted during research cruises aboard Nagasaki University’s training vessel Kakuyo-maru, using baited traps deployed on the seafloor (Anzai et al., 2026). After recovery, the isopods were transferred to seawater at approximately 10 °C and transported to the laboratory in insulated containers. In the laboratory, the isopods were maintained in a 200-L black circular holding tank under continuous darkness. The tank was equipped with aeration and a filtration system (Eheim 2260, Eheim, Germany), and the water temperature was maintained at 10 °C using a chiller (Cool Way BK210, Gex, Osaka, Japan). From this holding population, 29 individuals spanning a body-mass range of 1.7‒48.4 g were selected for metabolic measurements. To standardize their physiological condition and minimize the effects of recent feeding and specific dynamic action, these individuals were fasted for four weeks before respirometry. All experimental procedures were approved by the Animal Experiment Committee of the Faculty of Fisheries, Nagasaki University (approval no. NF-0095).

### 2.2 Measurement of oxygen consumption

Oxygen consumption rate 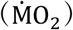 was measured using intermittent-flow respirometry at approximately 10 °C, following recommended procedures for aquatic respirometry (Killen et al., 2021) and the calculation approach previously applied to *B. doederleini* (Tanaka et al., 2023). Respiration chambers were selected according to specimen body mass, with chamber volumes ranging from 180 to 878 mL. The effective water volume of each chamber was determined from the mass of seawater contained in the chamber and corrected for the volume displaced by the experimental animal. The displaced volume was estimated from body mass using a tissue density of 1.225 g mL^-1^, as previously determined for *B. doederleini* (Tanaka et al., 2023).

Dissolved oxygen concentration was recorded continuously during each closed measurement period. Each closed measurement interval was defined from the onset of a consistent decline in dissolved oxygen concentration after chamber closure to the final data point before flushing caused oxygen concentration to increase. For each interval, the rate of change in dissolved oxygen concentration was estimated by linear regression against time. Background oxygen consumption was corrected by subtracting the slope measured in the corresponding empty chamber from that measured in the chamber containing the animal. For some measurements, a reliable background slope could not be obtained from the blank measurement conducted on the same day. In these cases, the blank slope measured on the closest available date using the same respiration chamber was used for correction.

Whole-animal oxygen consumption rate was calculated as:

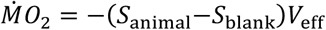

where *S*_animal_ is the slope of dissolved oxygen concentration measured in the chamber containing the animal (mg O_2_ L^-1^h^-1^), *S*_blank_ is the corresponding background slope measured in an empty chamber(mg O_2_ L^-1^h^-1^), and *V*_eff_ is the effective chamber volume after accounting for the volume displaced by the animal (L). The negative sign converts the decline in dissolved oxygen concentration into a positive oxygen consumption rate. Oxygen consumption rate was expressed as mg O_2_ L^-1^h^-1^.

### 2.3 Estimation of RMR, AMR and FAS

For each individual, three to six resting measurement cycles were conducted. To reduce the influence of handling and initial acclimation to the respiration chamber, only the final three resting cycles were considered for estimation of resting metabolic rate (RMR). Oxygen consumption was calculated separately for each cycle, and RMR was defined as the lowest background-corrected whole-animal oxygen consumption rate among the final three valid resting cycles. This procedure was based on the general approach of estimating resting or standard metabolic rate from the lowest stable oxygen uptake rates recorded after an initial acclimation period (Enders et al., 2006; Chabot et al., 2016).

To induce elevated metabolic demand, each individual was removed from the respiration chamber and subjected to 5 min of forced swimming at the same experimental temperature. Immediately after exercise, the individual was returned to the respiration chamber, and oxygen consumption was measured. Active metabolic rate (AMR) was defined as the background-corrected oxygen consumption rate obtained during the earliest post-exercise closed measurement interval. Accordingly, AMR was treated as the highest post-exercise oxygen consumption measured under the standardized protocol, rather than as a confirmed maximum metabolic rate. Factorial aerobic scope (FAS) was calculated for each individual as:

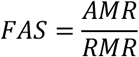

### 2.4 Statistical analyses

Body mass, RMR, AMR, and FAS were log10-transformed before analysis. Separate ordinary least-squares regressions were first used to describe the body-mass scaling relationships of RMR and AMR based on the allometric equation:

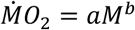

where 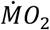 is whole-animal oxygen consumption rate, *M* is body mass in grams, *a* is the scaling coefficient, and *b* is the scaling exponent. The fitted relationships were back-transformed and expressed on the original measurement scale.

Because RMR and AMR were measured in the same individuals, their scaling relationships were compared using a linear mixed-effects model. Log10-transformed metabolic rate was modelled in relation to log10-transformed body mass, metabolic state (RMR or AMR), and their interaction, with individual identity included as a random intercept. The interaction term was used to test whether the scaling exponents of RMR and AMR differed. When the interaction was not significant, a reduced model without the interaction term was used to estimate a common scaling exponent.

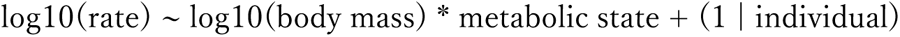

RMR was used as the reference metabolic state. The coefficient of log10 body mass therefore represented the RMR scaling exponent, whereas the AMR exponent was obtained as the sum of the body-mass coefficient and the body mass × metabolic state interaction. The interaction tested whether the AMR and RMR scaling exponents differed. Denominator degrees of freedom and p values were obtained using the Satterthwaite approximation, and state-specific slopes and their contrast were estimated using marginal trends. When the interaction was not significant, a reduced mixed-effects model without the interaction term was used to estimate a common scaling exponent.

Factorial aerobic scope was calculated as FAS = AMR/RMR. The relationship between FAS and body mass was tested by regressing log10(FAS) against log10(body mass). Because log10(FAS) = log10(AMR) - log10(RMR), the slope of this regression is algebraically equivalent to the difference between the AMR and RMR scaling exponents and therefore provided a paired confirmation of the mixed-model interaction. The average proportional elevation of AMR above RMR was summarized as the geometric mean of AMR/RMR with a 95% confidence interval.

To examine body-mass-independent covariation between RMR and AMR, residuals were obtained from their separate log-log regressions against body mass, and residual AMR was regressed against residual RMR. Potentially influential observations were identified using Cook’s distance, with values greater than 4/n treated as potentially influential. All 29 individuals were retained in the primary analyses, and sensitivity analyses were repeated after excluding potentially influential observations. The R^2^ values of the selected oxygen-decline intervals were retained as measurement-quality information but were not used as statistical exclusion criteria. Results are reported with 95% confidence intervals. A significance level of P < 0.05 was used for all statistical tests, which were performed using R (R Core Team, 2024, version 4.4.0).

## 3. Results

### 3.1 Body-mass scaling of RMR and AMR

**Resting metabolic rate and AMR, measured as post-exercise peak oxygen consumption, both increased significantly with body mass in Bathynomus doederleini (Fig. 1A, B). Separate ordinary least-squares regressions gave the following allometric relationships:**

**Fig. 1.**
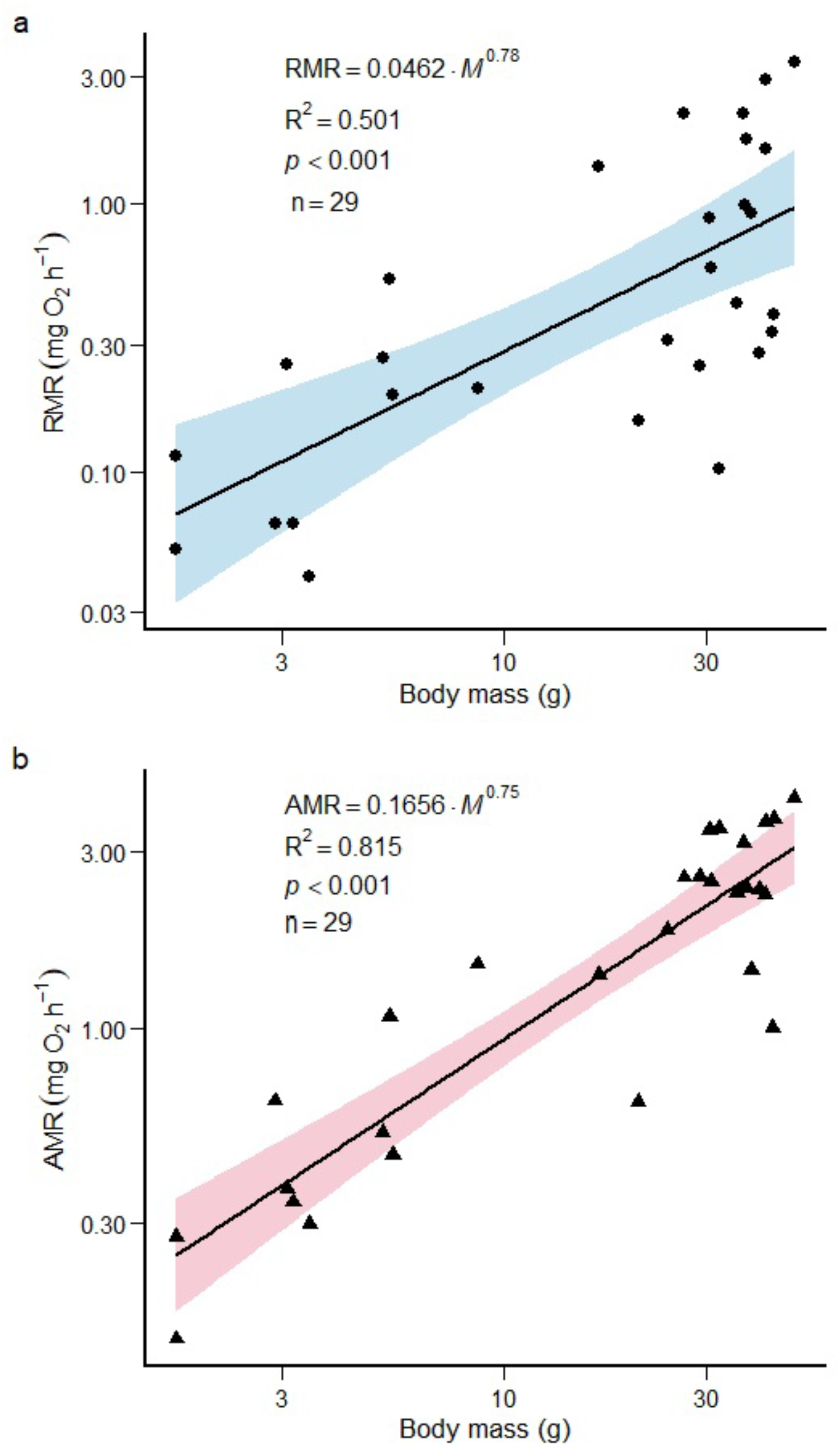
Ontogenetic scaling of resting metabolic rate (RMR) and active metabolic rate (AMR) in *Bathynomus doederleini*. (A) RMR increased significantly with body mass. (B) AMR, measured as post-exercise peak oxygen consumption, also increased significantly with body mass. Lines indicate fitted allometric regressions, and shaded areas indicate 95% confidence intervals.

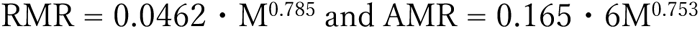

where M represents body mass in grams and metabolic rate is expressed as mg O2 h-1. The RMR exponent was 0.785 (95% confidence interval: 0.476-1.094; R^2^ = 0.501, p < 0.001), and the AMR exponent was 0.753 (95% confidence interval: 0.611-0.894; R^2^ = 0.815, p < 0.001). In the individual-level mixed-effects model, the body mass × metabolic state interaction was not significant (difference between AMR and RMR exponents = -0.032, 95% confidence interval: -0.330 to 0.267, p = 0.829). Thus, the data provided no evidence that RMR and AMR scaled with different exponents. A reduced common-slope model gave a common exponent of 0.769 (95% confidence interval: 0.580-0.957, p < 0.001). The geometric mean AMR/RMR ratio was 3.29 (95% confidence interval: 2.37-4.56), indicating that post-exercise oxygen consumption was, on average, approximately 3.3-fold higher than resting metabolism.

### 3.2 Size dependence of factorial aerobic scope

Consistent with the similar scaling of RMR and AMR, FAS showed no detectable relationship with body mass (R^2^ = 0.0018, p = 0.829; Fig. 2A). The fitted relationship was:

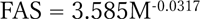

**Fig. 2.**
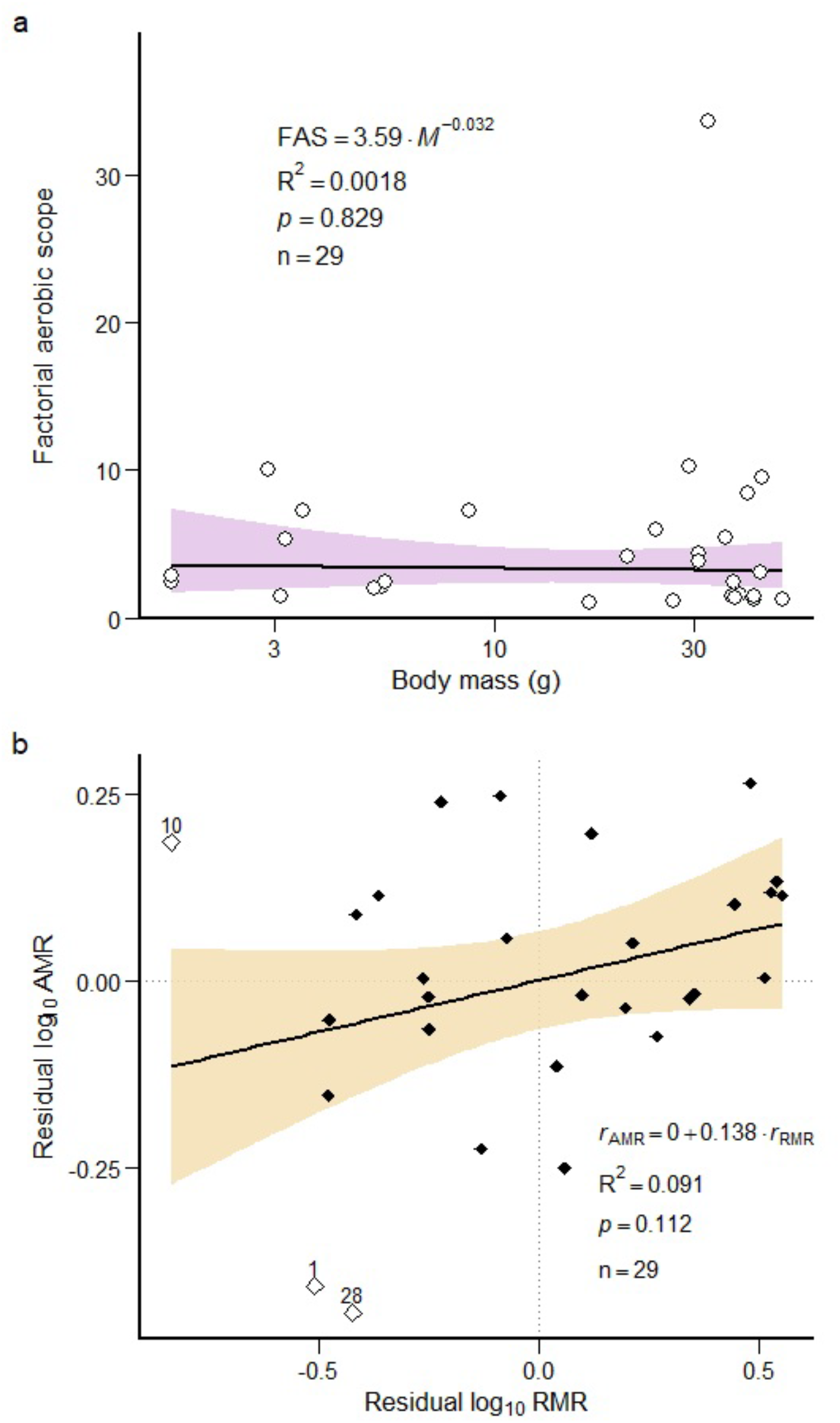
Size dependence of factorial aerobic scope and body-mass-corrected metabolic traits in *B. doederleini*. (A) Factorial aerobic scope (FAS = AMR/RMR) showed no significant relationship with body mass. (B) Residual AMR was not significantly related to residual RMR after adjusting for body mass. Shaded areas indicate 95% confidence intervals.

The FAS exponent was -0.0317 (95% confidence interval: -0.330 to 0.267). Median FAS was 2.83, with an interquartile range of 1.47-5.96. Although FAS varied substantially among individuals, this variation showed no directional trend with body mass.

### 3.3 Body-mass-corrected covariation between RMR and AMR

After accounting for body mass, residual RMR and residual AMR showed a weak positive relationship, but the association was not statistically significant (slope = 0.138, 95% confidence interval: -0.034 to 0.311, R^2^ = 0.091, p = 0.112, n = 29; Fig. 2B). Individuals with RMR values above those expected for their body mass therefore did not consistently exhibit AMR values above those expected for their body mass.

Cook’s distance identified three potentially influential observations (IDs 1, 10 and 28). After excluding these observations, the residual relationship remained non-significant (slope = 0.094, 95% confidence interval: -0.068 to 0.257, R^2^ = 0.056, p = 0.242, n = 26). The principal conclusion was therefore unchanged by the sensitivity analysis.

## 4. Discussion

### 4.1 Proportional scaling of resting and post-exercise metabolism

The principal finding of this study was that resting metabolic rate (RMR) and post-exercise peak oxygen consumption, treated here as active metabolic rate (AMR), increased approximately in parallel with body mass in *Bathynomus doederleini*. The scaling exponents of RMR and AMR did not differ significantly, yielding a common exponent of 0.77. In the common-slope model, AMR remained approximately 3.3-fold higher than RMR across the body-mass range examined. These results indicate that the activity-related aerobic response increased during growth in approximate proportion to resting maintenance demand.

The common exponent of 0.77 is close to the frequently discussed three-quarter-power scaling relationship. A comparative analysis of ontogenetic metabolic scaling in crustaceans reported a median exponent of 0.736 and a mean exponent of 0.735 ± 0.038 for benthic crustaceans (Glazier, 2006). Relative to these representative values, the common exponent of 0.77, and particularly the RMR exponent of 0.79, were numerically somewhat higher, suggesting that metabolic rate increases relatively steeply with body mass in the benthic *B. doederleini*. Nevertheless, the primary biological significance of the present findings lies not in their proximity to a particular theoretical or comparative value, but in the similarity of the RMR and AMR exponents. Metabolic scaling exponents can vary with taxonomic affiliation, lifestyle, activity level, physiological state, and environmental conditions (Glazier, 2005, 2009, 2010; Killen et al., 2010).

Resting and maximum or activity-related metabolic rates do not necessarily scale in parallel. In some aquatic ectotherms, differences between their scaling exponents produce ontogenetic changes in aerobic scope (Brett, 1965; Post and Lee, 1996; Killen et al., 2007; Zhang et al., 2014). When resting metabolism increases more steeply than activity-related metabolism, the proportional aerobic capacity available beyond maintenance may decline during growth. For example, in the deep-water New Zealand scampi *Metanephrops challengeri*, mass-specific SMR increased with body mass whereas mass-specific MMR decreased, suggesting that metabolic scope narrows in larger individuals (Heasman, 2022). However, that study analysed mass-specific metabolic rates and calculated metabolic scope from the difference between SMR and MMR, whereas the present study examined whole-animal metabolic rates and factorial aerobic scope.

Conversely, when activity-related metabolism increases more steeply than resting metabolism, larger individuals may acquire a greater proportional aerobic margin. Neither pattern was observed in *B. doederleini*. Although larger individuals had greater absolute oxygen demand, the proportional relationship between resting and post-exercise metabolism was maintained across the examined body-mass range. Thus, the increase in the metabolic cost of maintaining a larger body was accompanied by a corresponding increase in the capacity to elevate oxygen consumption after exercise. This coordinated scaling may help preserve the balance between maintenance demand and episodic activity throughout post-hatching growth.

### 4.2 Body-mass-independent factorial aerobic scope

Consistent with the similar scaling exponents of RMR and AMR, factorial aerobic scope (FAS = AMR/RMR) showed no significant relationship with body mass. Thus, proportional post-exercise aerobic capacity relative to maintenance metabolism was maintained across the body-mass range examined in *B. doederleini*.

Ontogenetic changes in aerobic scope may be associated with changes in locomotor mode, tissue composition, oxygen transport capacity, habitat use, and energy allocation (Post and Lee, 1996; Killen et al., 2007; Glazier, 2009). Species undergoing pronounced ecological or morphological transitions among larval, juvenile, and adult stages may consequently show substantial changes in the relative scaling of resting and maximum metabolism. In contrast, *B. doederleini* appears to retain a broadly benthic scavenging lifestyle throughout the post-hatching body mass range examined here (Soong and Mok, 1994). This continuity in ecological role may contribute to the stable balance between maintenance and activity-related metabolism during growth.

Although FAS showed no directional relationship with body mass, substantial variation was observed among individuals. Individual variation in metabolic rate is common and can be repeatable, although its magnitude and ecological significance vary among metabolic traits and taxa (Nespolo and Franco, 2007; Careau et al., 2008; Norin and Malte, 2011). The broad variation in FAS therefore indicates that the aerobic metabolic phenotype of *B. doederleini* is not determined by body mass alone. Physiological condition, behaviour, molt stage, nutritional history, and responsiveness to exercise may also contribute to variation among individuals.

### 4.3 Individual-level metabolic covariation and ecological implications for a deep-sea scavenger

Although RMR and AMR scaled proportionally at the population level, no significant relationship was detected between the residuals obtained from their respective regressions against body mass. Individuals with RMR values above those expected for their body mass did not consistently exhibit AMR values above those expected for their body mass. Studies of fishes have likewise reported inconsistent relationships between mass-corrected resting and maximum or activity-related metabolism, with outcomes depending on the metabolic traits and analytical level considered (Norin and Malte, 2011, 2012; Auer et al., 2017; Reemeyer and Rees, 2020).

In *B. doederleini*, RMR may primarily reflect the energetic costs of maintaining cells, organs, and tissues, whereas post-exercise oxygen consumption may additionally depend on exercise intensity, behavioral responsiveness, anaerobic energy use, and recovery processes. These partly distinct determinants may explain the weak relationship between the two traits after accounting for body mass. Accordingly, the present results provided no clear evidence that body-mass-independent FAS arose from a strong, fixed individual-level relationship in which individuals with high RMR invariably possessed proportionally high AMR.

This individual-level variation can be interpreted in the context of the spatially and temporally heterogeneous food supply of the deep sea. Large organic falls can create intense but localized resource pulses, whereas the intervals between such events may be characterized by limited food availability (Dayton and Hessler, 1972; Smith and Baco, 2003). A high RMR imposes a continuous energetic cost, whereas high activity-related aerobic capacity may provide benefits during restricted periods of movement toward food, feeding, digestion, and post-activity recovery. The ecological value of a particular metabolic phenotype may therefore depend on food availability and opportunities for activity (Auer et al., 2015, 2017).

Metabolic patterns in deep-sea animals are shaped not only by physical conditions such as low temperature, high hydrostatic pressure, and oxygen availability, but also by ecological energy demand and lifestyle (Childress, 1995; Seibel and Drazen, 2007; Yagi et al., 2025). Crustaceans inhabiting food-limited environments can reduce locomotory, ventilatory, and respiratory activity during prolonged starvation (Hervant et al., 1997). In *B. doederleini*, feeding and temperature have also been shown to influence metabolic rate, indicating that metabolism responds to both nutritional and environmental conditions (Tanaka et al., 2023). Furthermore, low basal metabolic rates and prolonged food retention have been associated with extreme starvation tolerance in deep-sea giant isopods, emphasizing the potential importance of reduced maintenance expenditure under unpredictable food supply (Yuan et al., 2026).

Taken together, the proportional scaling of RMR and AMR in *B. doederleini* indicates that activity-related aerobic capacity relative to maintenance demand remained broadly similar across the body-mass range examined. At the same time, the weak and statistically non-significant relationship between their residuals indicates that strong individual-level covariation was not clearly supported by the present sample. Direct measurements of spontaneous activity, food-search behaviour, and feeding performance will be required to determine whether variation in metabolic phenotype has ecological consequences under episodic food availability.

### 4.4 Conclusions

In *Bathynomus doederleini*, RMR and AMR scaled proportionally with body mass, resulting in an exercise-induced FAS that was independent of body size. Thus, although absolute metabolic demand increased during growth, activity-related aerobic capacity relative to maintenance metabolism was maintained.

In contrast, no significant relationship was detected between the body-mass regression residuals of RMR and AMR. Parallel population-level scaling therefore did not necessarily reflect strong individual-level coupling between the two metabolic traits. These findings emphasize the importance of distinguishing average body-mass-related changes from individual variation in metabolic phenotype when evaluating aerobic performance in deep-sea scavengers.

## Acknowledgements

We thank the crew and staff who assisted with animal collection and maintenance.

## Ethics statement

The animals used in this study were marine invertebrates, and no specific animal ethics approval was required under the applicable institutional guidelines.

## Author contributions

S.T. and M.Y. designed the study. S.T., M.Y., S.I., and S.A. collected or maintained animals and contributed to data acquisition. S.T. and M.Y. analysed the data and drafted the manuscript. All authors reviewed and approved the final manuscript.

## Data availability

Data will be made available upon reasonable request.

